# Structure and function of a neocortical synapse

**DOI:** 10.1101/2019.12.13.875971

**Authors:** Simone Holler-Rickauer, German Köstinger, Kevan A.C. Martin, Gregor F.P. Schuhknecht, Ken J. Stratford

## Abstract

Thirty-four years since the small nervous system of the nematode *C. elegans* was manually reconstructed in the electron microscope (EM)^1^, ‘high-throughput’ EM techniques now enable the dense reconstruction of neural circuits within increasingly large brain volumes at synaptic resolution^2–6^. As with *C. elegans*, however, a key limitation for inferring brain function from neuronal wiring diagrams is that it remains unknown how the structure of a synapse seen in EM relates to its physiological transmission strength. Here, we related structure and function of the same synapses to bridge this gap: we combined paired whole-cell recordings of synaptically connected pyramidal neurons in slices of mouse somatosensory cortex with correlated light microscopy and high-resolution EM of all putative synaptic contacts between the neurons. We discovered a linear relationship between synapse size (postsynaptic density area) and synapse strength (excitatory postsynaptic potential amplitude), which provides an experimental foundation for assigning the actual physiological weights to synaptic connections seen in the EM. Furthermore, quantal analysis revealed that the number of vesicle release sites exceeded the number of anatomical synapses formed by a connection by a factor of at least 2.6, which challenges the current understanding of synaptic release in neocortex and suggests that neocortical synapses operate with multivesicular release, like hippocampal synapses^7–11^. Thus, neocortical synapses are more complex computational devices and may modulate their strength more flexibly than previously thought, with the corollary that the canonical neocortical microcircuitry possesses significantly higher computational power than estimated by current models.

## Introduction

Since the pioneering studies of the neuromuscular junction by Katz and colleagues^12^, it has been known that synaptic transmission is probabilistic and can be described by simple binomial models with three parameters: the number of release sites, quantal size, and release probability^13^. For some central synapses, these parameters have been linked to synaptic ultrastructure: in hippocampus, the number of membrane-bound vesicles in the readily-releasable pool scales with the size of the presynaptic active zone (the membrane region where vesicles are docked and released), and both correlate with release probability^14–18^. Postsynaptically, the estimated number of AMPAr (α-amino-3-hydroxy-5-methyl-4-isoxazolepropionic acid receptors) is proportional to postsynaptic density (PSD) area in hippocampal and cerebellar synapses^19,20^. The absolute strength of synaptic transmission can be measured physiologically as the amplitude of the excitatory postsynaptic potential (EPSP), which is the product of all three quantal parameters in binomial statistics, but it remains experimentally untested for any synapse in the brain if and how the EPSP amplitude relates to PSD area. A major limitation towards assessing this important size-strength relationship in neocortex from the previous structure-function observations is that it still remains controversial how many release sites exist at single synapses^21–26^.

The prevailing one-synapse, one-quantum hypothesis suggests that each synapse contains only a single site at which neurotransmitter is released probabilistically. This conclusion arises from the finding that the number of putative synaptic contacts between connected pyramidal neurons – estimated principally from light microscopic (LM) observations – is on average not significantly different to the number of release sites estimated from quantal analysis^21,23^. This implies that the entire synaptic receptor array must be effectively saturated by the release of a single vesicle^27–29^. While the one-synapse, one-quantum hypothesis has been one of the critical factors in describing the principles of neocortical wiring^30–33^, the critical caveat is that a putative synapse (the close physical proximity of axon and dendrite in LM) is not in fact predictive for the existence of an actual synapse, which can be identified only in the EM^2,4,6^. The existence of synapses at LM appositions was only checked with EM for a subset of experiments in these studies^21,23^ and even in these cases, reliable synapse identification was compromised by the electron-dense biocytin reaction product in synaptic terminals^21^.

Here, we set out to resolve if neocortical synapses contain only a single or multiple release sites and to establish the precise nature of the relationship between anatomical synapse size and physiological transmission strength. We recorded synaptic transmission between pairs of connected layer 2/3 (L2/3) pyramidal neurons in slices of mouse somatosensory cortex at postnatal days 21 to 30 and measured mean EPSP amplitude and variance. During recordings, the cells were filled with biocytin, which allowed us to identify all potential synaptic contacts between the axon of the presynaptic neuron and the dendrites of the postsynaptic neuron in the LM. We then recovered and examined every single LM contact in the EM to identify all sites at which actual synapses were formed between the recorded cells. Finally, the high-fidelity ultrastructural preservation of the tissue in combination with a series of high-magnification serial-section EM procedures enabled us to measure the PSD areas of all identified synapses despite biocytin reaction product in the synaptic terminals.

## Results

Of 59 recorded pairs, only 10 pairs passed both our stringent electrophysiological and anatomical quality controls and formed the final dataset (see *Methods*). We required mean and variance of EPSP recordings to remain stable for ≥ 100 consecutive trials and for the dendrites and axons to be entirely filled with biocytin, as assessed during the LM reconstructions. Unitary EPSP amplitudes during stable epochs of recording ranged from 0.15 mV to 2.25 mV (mean ± standard deviation: 1.05 ± 0.70 mV).

In LM, we identified 40 axodendritic appositions between the 10 pairs (range: 1 to 7, mean ± standard deviation: 4.0 ± 1.8), consistent with previous studies in rodents^21,23,31^. Following slice recordings, we achieved high-fidelity ultrastructural preservation and were able to resolve synapse specializations, including the PSD, the synaptic cleft, and vesicles in the presynaptic bouton (Figs. 1, 2, extended Fig. 1, 2). Serial-section EM of all 40 LM appositions revealed conclusively that synapses between presynaptic axon and postsynaptic dendrite were formed at only 16 appositions (mean: 1.6). Of the 10 pairs, 6 were connected by 2 synapses, and 4 by a single synapse (extended Fig. 3). In all cases, synapses could be identified unambiguously: axodendritic contacts extended over multiple sections and the hallmarks of synaptic specializations were present. Importantly, we could always detect when no synapse was formed between the labeled presynaptic axon and postsynaptic dendrite at identified LM appositions: in all such cases, EM revealed a distinct gap between the labeled structures and no synaptic specializations. Furthermore, the labeled presynaptic boutons and dendritic spines at these contacts were found in the EM to form synapses with unlabeled partners in the neuropil and not with the labeled dendrites and axons of the other neuron, respectively (Figs 1 f, 2). Notably, we were not able to discover any LM features that would distinguish between synapse forming and non-synapse forming appositions, which is in accordance with previous studies^2,4^

**Figure 1.**
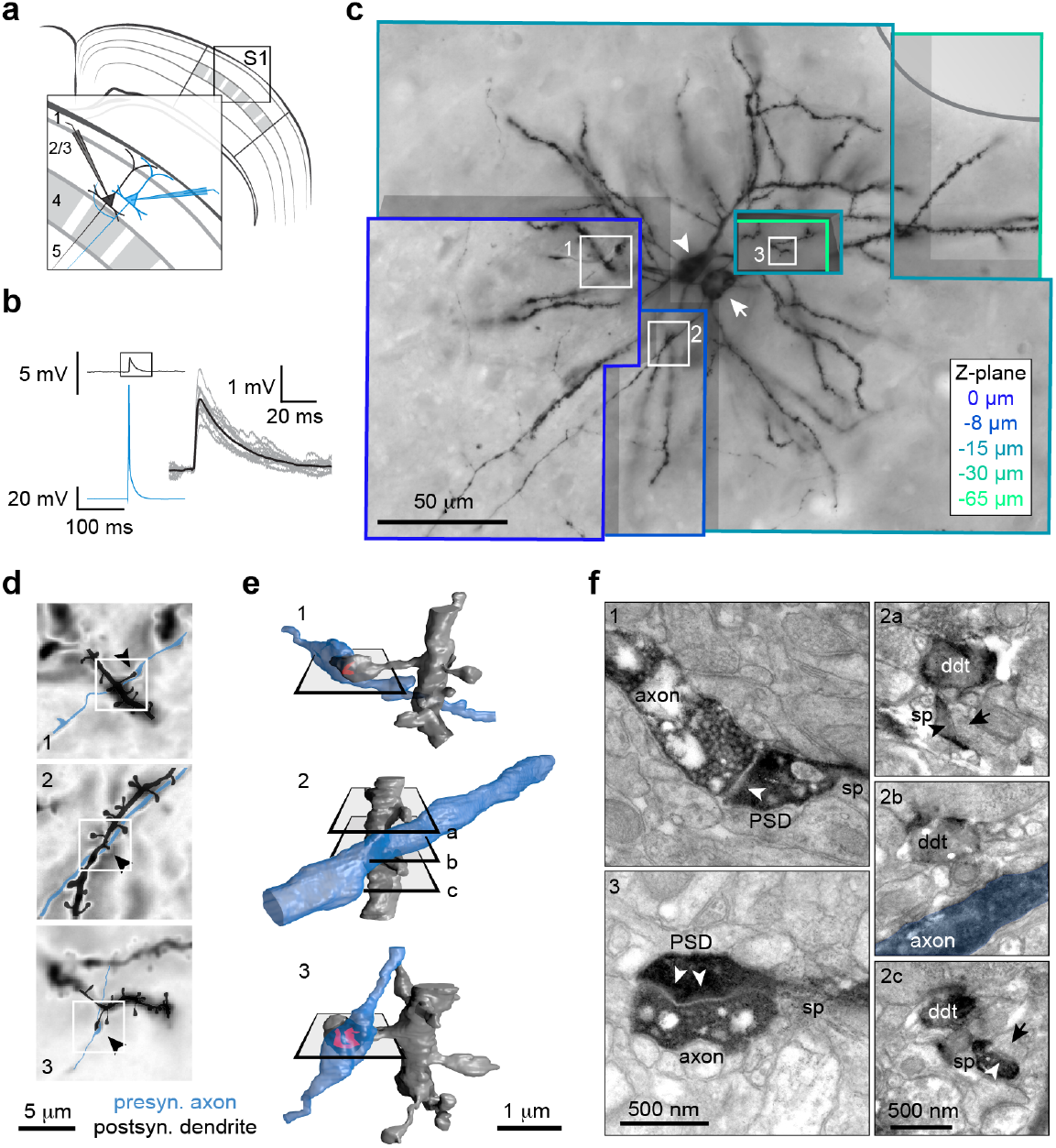
Relating electrophysiology and ultrastructure of synapses between L2/3 pyramidal neurons in mouse barrel cortex (S1). **a** Schematic of whole-cell recordings of synaptically connected L2/3 pyramidal neurons in slices of sensory cortex. **b** Left, averaged traces of evoked action potential in the presynaptic neuron (blue) and resulting EPSP in the postsynaptic neuron (black). Right, enlarged waveforms of averaged EPSP (black) and 10 randomly-selected single-trial EPSPs (grey). **c** 3D LM stack containing the biocytin-reacted neurons after paired recordings shown in b. Three appositions (numbered boxes) were identified between presynaptic axon and postsynaptic dendrite. Arrow, soma of presynaptic neuron; arrowhead, soma of postsynaptic neuron; different z-planes through the stack are color-coded and normalized to topmost section; grey line, cortical surface. **d** High-magnification LM of appositions (numbering corresponds to c) overlaid with drawings of contact points made from LM stacks and EM reconstructions (not shown). Arrowheads, locations of potential synapses. White boxes indicate positions of EM reconstructions in e. **e** 3D EM reconstructions revealed that synapses between presynaptic axon and postsynaptic dendrite were formed only at appositions 1 and 3. Red, PSD; planes indicate positions of electron micrographs in f. **f** Electron micrographs of cross-sections through appositions, as indicated by planes in e. The presynaptic axon formed synapses on dendritic spines (sp) that emerged from postsynaptic dendrites (ddt) at appositions 1 and 3. Presynaptic vesicles, synaptic cleft, and PSD (arrowhead) could be distinguished despite biocytin reaction product. At apposition 2, the postsynaptic dendritic spines formed synapses (arrowheads) with unlabeled boutons (arrows). At the point of closest proximity, a physical separation remained between presynaptic axon and postsynaptic dendrite (2b).

**Figure 2.**
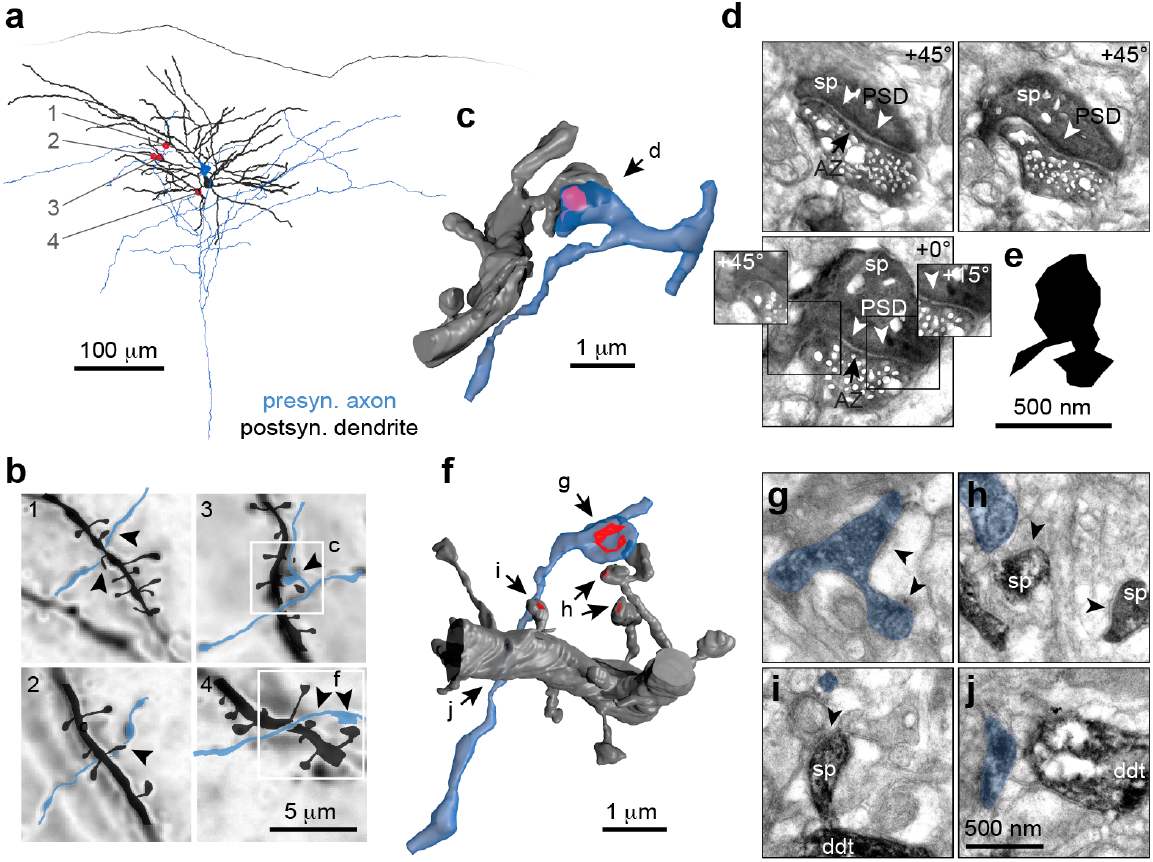
At most LM appositions, no synapses are formed between the recorded pyramidal neurons. **a** Full 3D LM reconstruction of presynaptic axon and postsynaptic dendrite of synaptically connected pyramidal neurons (somata and cortical surface shown, presynaptic dendrite and postsynaptic axon excluded for clarity). Four appositions (red dots, numbered) were identified. **b** High-magnification LM of appositions (numbering corresponds to a), overlaid with drawings of contact points made from LM stacks and EM reconstructions (not shown). Arrowheads, locations of potential synapses; white boxes, positions of EM reconstructions in c, f. At appositions 1 and 2, EM revealed that no synapses were formed between presynaptic axon and postsynaptic dendrite (not shown). **c** EM reconstruction of apposition 3 revealed that a synapse was formed between the presynaptic axon and a spine emerging from the postsynaptic dendrite. Arrow, position of micrographs shown in d; red, PSD. **d** Electron micrographs of consecutive cross-sections through synapse in c. Presynaptic vesicles, active zone (AZ; arrow) and PSD (arrowheads) are visible despite biocytin reaction product; sp, spine. Tilt angles, at which micrographs were acquired, are indicated at top right. Insets show additional tilts of regions indicated by boxes. **e** 2D projection of fully reconstructed PSD. **f** EM reconstruction of apposition 4 revealed that no synapses were formed between presynaptic axon and postsynaptic dendrite at 4 potential synaptic locations (arrows); red, PSDs formed with unlabeled structures in the neuropil. **g – j** Electron micrographs of locations indicated by arrows in f. Blue, presynaptic axon; arrowheads, synaptic specializations; sp, dendritic spine; ddt, dendritic shaft. **g** Presynaptic axon forming perforated synapse with unlabeled dendritic spine. **h** Postsynaptic dendritic spines forming synapses with unlabeled axons and not the labeled presynaptic axon. **i** Postsynaptic spine forming synapse with unlabeled axon. Presynaptic axon traversing section. **j** Presynaptic axon traversing in close proximity to postsynaptic dendrite without forming a synaptic bouton.

We employed a battery of technical procedures that allowed us to measure the PSD areas of the 16 identified synapses despite biocytin reaction product in the dendritic spines (see *Methods*). Briefly, we tilted all ultrathin sections containing synapses at six angles to obtain an optimal perpendicular imaging plane through the synaptic cleft (Fig. 2, extended Fig. 1). Each synapse was reconstructed independently by two experienced practitioners who were blind to the electrophysiological results. As a cross-check, we then validated the reliability of our PSD reconstructions against the known linear relationship between PSD area and spine head volume^34,35^: we found the same PSD-spine head relationship for biocytin-labeled synapses as for a ‘ground-truth’ dataset that we acquired from unlabeled synapses in adjacent neuropil (Fig. 3 a).

**Figure 3.**
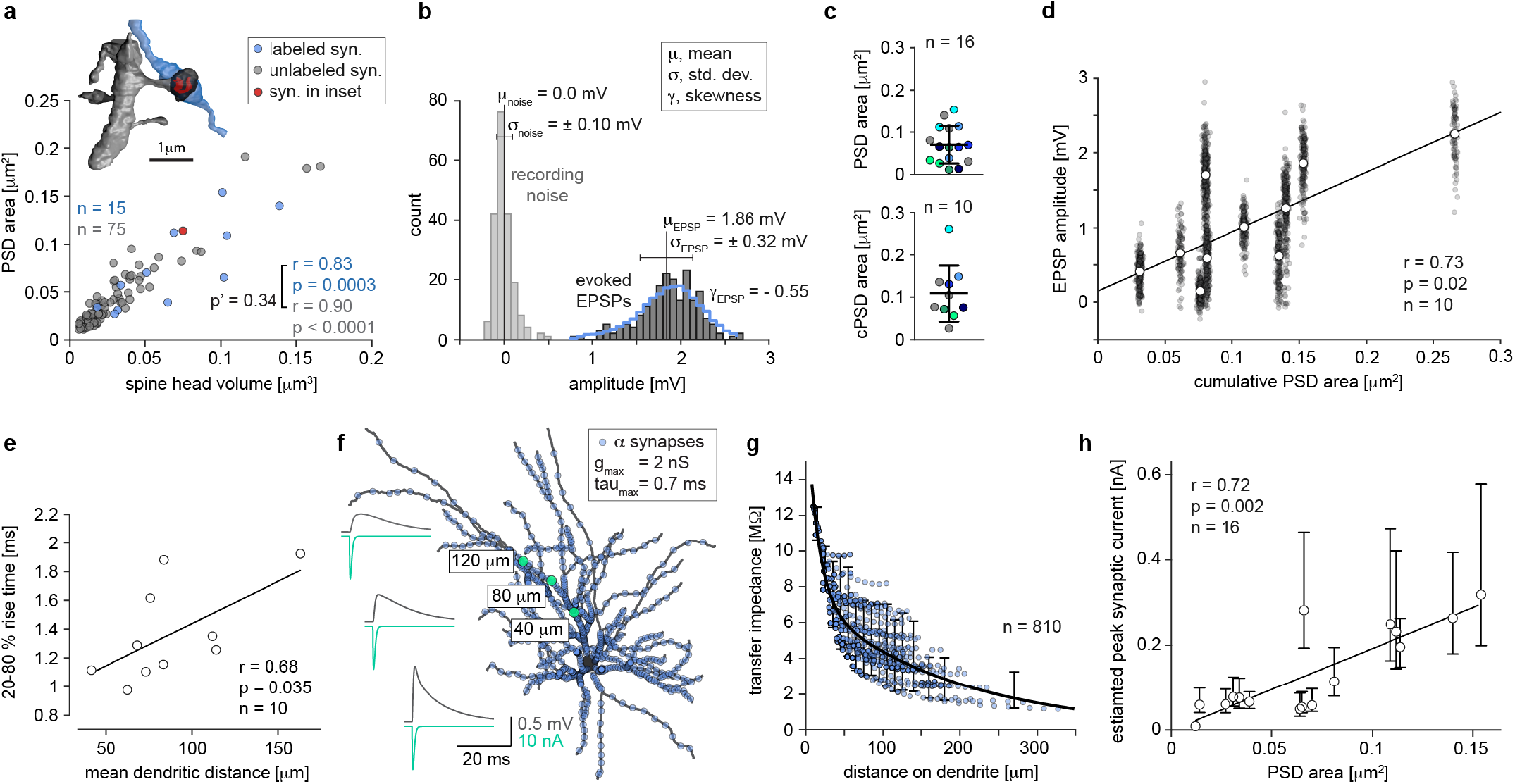
Synapse size predicts synaptic transmission strength. **a** Correlation between spine head volume and PSD area for biocytin-filled synapses and unlabeled, naïve synapses reconstructed from the same neuropil; p’, p value of Fisher-Z-transformation to test differences between correlation coefficients. One biocytin-filled synapse excluded because it was formed on a dendritic shaft. Inset, EM reconstruction of representative biocytin-filled synapse (synapse 3 in Fig. 1, rotated for clear view on spine head); blue, axon; grey, dendrite; black, spine head; red, PSD. **b** Evoked EPSP amplitudes and recording noise during stable epoch of recording (n = 200 trials) for the experiment shown in Fig. 1. 7-point moving average (blue) overlaid over EPSP distribution to highlight its left-skew (i.e. negative skewness). **c** Top, distribution of PSD areas of all 16 synapses between the 10 connected pairs. Synapses made by the same connection indicated by corresponding colors; grey, connections with single synapses. Bottom, distribution of cumulative PSD (cPSD) areas of 10 pairs; corresponding color-code; horizontal line, mean; error bars, standard deviation. **d** Relationship of mean EPSP amplitude (white dots) with cumulative PSD area for the 10 connected pairs. Grey points, amplitudes of all evoked EPSPs during stable epochs across experiments. Mean EPSP amplitudes were fit with a line (slope = 7.97 ± 2.44). **e** EPSP rise-time correlates with mean dendritic distance of identified synapses to soma. **f** Compartmental model from volumetric reconstruction of recorded pyramidal neuron. 810 alpha synapses were simulated along entire dendritic tree. Insets, time courses of synaptic currents (green) at 3 synaptic sites on same dendrite (green dots) and the resulting somatic EPSPs (grey); g_max_, peak synaptic conductance; tau_max_, time constant of synaptic conductance. **g** Transfer impedance for all simulated synapses as a function of synaptic distance to soma, fitted with double exponential function; errors bars, 2.5 - 97.5 percentiles for the transfer impedance in bins of dendritic distance (see *Methods*). **h** Relationship between PSD area and estimated peak synaptic current amplitude at the synaptic locations for all 16 synapses found in the study; 95% confidence intervals indicated (constructed from percentiles in g). Data was fit with a line (slope = 1.93 ± 0.34). r = non-parametric Spearman correlation coefficient; lines were fit using linear regression.

We computed the cumulative PSD area, i.e. the total PSD area for a connection, by summing the individual PSD areas for those connections with two synapses. We found that the cumulative PSD area was positively correlated with the mean EPSP amplitude of connections (Fig. 3 d). Given the characteristically large trial-to-trial variability of evoked EPSP amplitudes and the restricted dataset size resulting from our stringent quality criteria, the question arose as to the robustness of the PSD-EPSP correlation. To address this, we generated new datasets of mean EPSP amplitudes that were generated from bootstrap-resampled EPSP amplitude distributions and computed their correlation with the cumulative PSD area (n = 10,000). All of these resampled correlations were significant (not shown), suggesting that the synaptic size-strength relationship is robust when at least 100 evoked responses are averaged. Additionally, we computed correlations between randomly-selected single-trial EPSPs and cumulative PSD area and found that only 60% of these correlations were significant (n = 10,000; not shown). Thus, while the size-strength relationship in our dataset of 10 synapses is weak for single trials, it becomes statistically very robust for a large dataset of EPSP recordings.

Somatic EPSP recordings give only a limited picture of EPSPs originating in the dendrites, because input impedance changes along the dendritic tree and because voltage attenuation increases as a function of synaptic distance to the soma^36^. The 16 synapses in this study were located at mean distances from the soma of 89 ± 45 μm (range: 33 μm to 235 μm). In search of a footprint for distance-dependent effects on our EPSPs, we found that EPSP rise-times indeed correlated with mean synaptic distance from soma (Fig. 3 e). Therefore, we additionally estimated the peak current evoked at each synapse (I_syn_) as a metric for transmission strength that is independent of input impedance and voltage attenuation. First, we generated a compartmental model of a representative L2/3 pyramidal neuron and computed the dendro-somatic transfer impedance, which is a measure for the transfer efficiency of synaptic signals to the soma and is defined as the ratio of somatic EPSP amplitude in response to the peak current injected at the location of the synapse^36–38^ (Fig. 3 f). The transfer impedance as a function of the dendritic distance of synapses to the soma was well-described by a double-exponential function (Fig. 3 g). We then approximated cumulative peak synaptic current for each of our 10 connections and found that it correlated with the cumulative PSD area (r = 0.65, p = 0.049, not shown). We sought further to relate physiological transmission strength to anatomical synapse size for each of the 16 synapse that were recorded in this study. Therefore, we approximated the peak synaptic current for each of these 16 synapses under the assumption that if two synapses (A and B) were formed by a connection, the ratio of PSD areas (PSD^A^ to PSD^B^) would be reflected in the ratio of synaptic currents (I_syn_^A^ to I_syn_^B^) and that the two synaptic potentials summate linearly at the soma (extended Fig. 4). In addition, we computed 95% confidence bounds for the synaptic currents. Importantly, we found that the synaptic currents that we estimated in this manner correlated with the PSD areas of the respective synapses (Fig. 3 h).

To test whether our recorded synapses contained only a single or multiple release sites, we derived the quantal parameters from our electrophysiological data using quantal analysis and investigated their anatomical substrate. Importantly, the mode of vesicle release should critically determine the structure-function relationship of both the number of release sites and the quantal size. If the one-synapse, one-quantum hypothesis applies, the number of anatomical synapses found in EM should equal the number of release sites. Because AMPAr are thought to approach full occupancy upon single-vesicle release^27,29^, it follows that PSD area (proportional to the total number of AMPAr in the receptor array) should correlate with the quantal size (the electrical effect of a single quantum). Conversely, if synapses operate with multivesicular release, the number of release sites would exceed the number of synapses. Furthermore, since multiple quantal release events within the same synapse must lead to increasing numbers of occupied AMPAr, a correlation between quantal size and PSD area would not be expected.

Because our stringent quality criteria excluded the majority of experiments from structure-function analyses, we required a simple and reliable method of quantal analysis that did not place additional prohibitive constraints on our physiological recordings. Thus, we could not rely on mean-variance analysis, which requires varying the recording conditions^21^, or the method of fitting binomial models to histograms with equally-spaced peaks, which is applicable only on selected subsets of experiments where ‘peaky’ amplitude histograms occur^23^. Instead, we built upon existing statistical methods^39,40^ and developed and validated a novel form of quantal analysis, which we term *Statistical Moments Analysis of Quanta* (SMAQ). Under the common assumption that quantal transmission obeys simple binomial statistics^23,25^, the mean^40^, standard deviation^40^, and skewness^39^ of experimentally recorded EPSP distributions can be expressed as functions of the number or release sites, release probability, and quantal size. Because skewness is independent of quantal size, unique analytical solutions for all quantal parameters can be obtained if the first three statistical moments of EPSP distributions are determined experimentally. Quantal analysis techniques rely on statistical analyses of relatively small datasets that are subject to significant biological and experimental noise. When confidence intervals are computed for estimated quantal parameters^23^, they are large, indicating that quantal analysis is generally imprecise and subject to significant uncertainty. To assess SMAQ reliability, we derived 95% confidence intervals for the estimated quantal parameters (see *Methods*). These confidence intervals were inspired by Bayesian logic and proved to be more rigorous (i.e. they produced wider confidence bounds) when compared to confidence intervals derived from classical bootstrap-resampling techniques^23^ (extended Fig. 5). As a cross-check of the validity of SMAQ, we were able to exploit 5 recorded EPSP histograms that had equally-spaced peaks (indicating discrete multiples of the quantal size), but that had failed our anatomical quality criteria. Analyzing these experiments both with SMAQ and by the conventional method of fitting binomial models to the histograms revealed that none of the solutions differed significantly between methods for the number of release sites, release probability, and quantal size. Likewise, comparing confidence intervals derived by bootstrap-resampling between methods suggested that SMAQ and fitting binomial models to histograms achieve similar precision (extended Fig. 5).

We used SMAQ to recover the number or release sites (mean ± standard deviation: 6.9 ± 4.2), release probability (0.6 ± 0.2), and quantal size (0.28 ± 0.15 mV) for the 10 connections and computed Bayesian-inspired confidence intervals for our predictions (Fig. 4 a, b, d). These values are in excellent agreement with previous quantal parameter estimates in rodent neocortex using different quantal analysis methods^21,23,25,41,42^. We found that the PSD areas of the 16 synapses correlated with the release probability of the connection (r = 0.62, p = 0.012) (Fig. 4 c), which was assumed to be uniform between all release sites of a connection in accordance with a simple binomial model. PSD area is known to match exactly the area of the active zone^14,15,25^ and thus, we show that the known correlation between active zone area and release probability in hippocampal synapses^17,18^ applies also to neocortical synapses. Critically, our estimates for the number or release sites exceeded the number of anatomical synapses for all connections, on average by a factor of 4.3. More importantly, the Bayesian-inspired, lower confidence interval bounds for the estimated number or release sites exceeded the number of anatomical synapses in all experiments but one (experiment 9 in Fig. 4 d) – on average by a factor of 2.6. This is a significant result that can only be reconciled with the notion that single neocortical synapses contain multiple release sites and that multivesicular release is the predominant mode of vesicle release between L2/3 pyramids. In accordance with this, we found no correlation between the estimated quantal size and the PSD areas of the 16 synapses in the study (p = 0.072; quantal size was assumed to be uniform when two synapses were formed by a connection in accordance with a simple binomial model).

**Figure 4.**
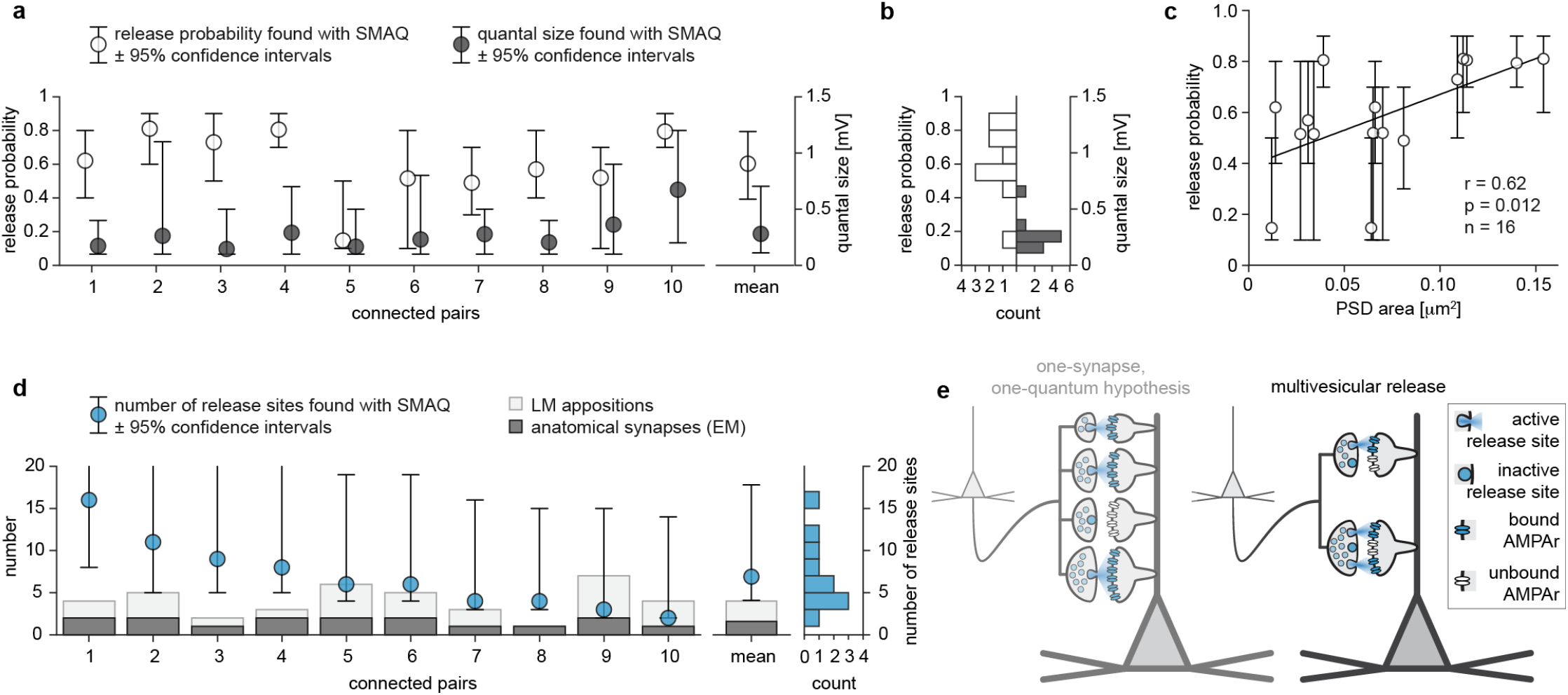
Synapses between L2/3 pyramidal neurons are capable of multivesicular release. **a** SMAQ solutions for release probability (left y axis) and quantal size (right y axis) with corresponding 95 % confidence intervals for the 10 connected pairs. Pairs are sorted by decreasing number of release sites, as shown in d. **b** Histograms of SMAQ solutions for release probability (left) and quantal size (right) plotted on same y-axes as a. **c** Release probability correlates with PSD area. r = non-parametric Spearman correlation coefficient; line was fit with linear regression; slope = 2.81 ± 1.04). **d** Left, SMAQ solutions for the number of release sites with corresponding 95 % confidence intervals plotted with the respective number of LM appositions and synapses found in EM for the 10 connected pairs. Right, histogram of SMAQ solutions for the number of release sites. **e** Schematic showing the two contrasting hypotheses of release mode at neocortical synapses. Left, according to the one-synapse, one-quantum hypothesis, each synapse contains a single release site and release of one vesicle saturates all postsynaptic AMPAr; this model is inconsistent with our findings. Right, our data support the model of multivesicular release, in which single neocortical synapses contain multiple release sites and release of a single vesicle is insufficient to saturate all postsynaptic AMPAr.

## Discussion

Because previous studies have estimated putative synapse numbers from LM with supporting data from less stringent EM^21,23,30,31,43^, they have supported the prevailing dogma that excitatory neocortical synapses contain only a single release site^21,23^. While the existence of a physical axodendritic apposition is a prerequisite for synapse formation, our high-fidelity correlated LM-EM for every single LM apposition revealed that synapses between pre- and postsynaptic neuron are formed at only a fraction of LM appositions, which is supported by ultrastructural analyses of neocortical and hippocampal neuropil and theoretical work on general principles of neocortical connectivity^2,4,6,44–46^. Our observation that multivesicular release occurs at synapses that span one order of magnitude in size is consistent with biophysical models suggesting that the highly localized potency of glutamate to activate AMPAr permits multiple release events within the same synapse^9,47^. Importantly, experimental studies and simulations have provided overwhelming evidence that also hippocampal synapses are capable of releasing multiple synaptic vesicles simultaneously^8–11^, which raises the intriguing hypothesis that multivesicular release is in fact a cardinal feature of excitatory synapses in the cerebral cortex. Multivesicular release at neocortical synapses means, however, that the number of release sites cannot be obtained by simply counting either the number of LM contacts or the number of synapses in EM. Likewise, the number of anatomical synapses cannot be derived from electrophysiological recordings.

At the same time, fundamental implications for neocortical computation arise: firstly, synapses operating with multivesicular release constitute more complex computational devices and likely endow the neocortical circuitry with significantly higher computational power than previously supposed under the one-synapse, one-quantum hypothesis, particularly if individual release sites at a synapse were found to be independently regulated. Secondly, multivesicular release enables plastic tuning of synaptic efficacies by adjusting the number of release sites within synapses without need for structural remodeling. This likely occurs in hippocampus, where the number of identified nanodomains per synapse scales with PSD area^11^. Thirdly, estimates of the number of presynaptic neurons that form synapses with a single neocortical pyramidal neuron have been based on the ratio of dendritic spine counts and the average number of putative synapses per connection^30,31^. Because synapses are formed at only a minority of putative synaptic locations, these studies have likely underestimated the actual presynaptic convergence in neocortex^32,33^. Each pyramidal neuron is connected to more presynaptic partners than estimated, which endows the neocortical circuitry with a higher computational capacity.

While the exact structural basis of the release site remains unresolved, a promising candidate are trans-synaptic nanocolumns as discovered in hippocampus^11,48,49^, whose putative role as release sites would be testable by employing super-resolution techniques to neocortical synapses. Intriguingly, our prediction that at least 2.6 release sites should be contained within single neocortical synapses agrees well with the 2.4 to 2.5 nanodomains that are found on average per hippocampal synapse^11^.

Finally, we have now provided first direct experimental evidence for a linear relationship between synapse size and synaptic transmission strength, which supports the general functional relevance of quantifying neocortical synapse size through EM. Intriguingly, we found a robust size-strength dependency only when many responses were temporally averaged in the very stable recordings we achieved in slices. The stochastic nature of synaptic transmission and the resulting large trial-to-trial variability in synaptic responses could weaken the observed PSD-EPSP correlation *in vivo*. However, the robustness of the size-strength relationship that we found for only 10 averaged EPSP distributions suggests that the dendritic tree of neocortical pyramidal neurons, which forms 10^3^ – 10^4^ synapses, could serve to maintain the observed PSD-EPSP correlation on a trial-to-trial basis *in vivo* by spatial averaging across its large synapse population. Thus, the linear relationship we established between synapse size and synaptic transmission strength provides the experimental means for extending the simple binary label of ‘connected’ or ‘not-connected’ in neocortical wiring diagrams to assigning actual physiological weights to synaptic connections. This is a key step on the path towards simulating information flow within neocortical connectomes.

## Extended Figures

**Extended figure 1.**
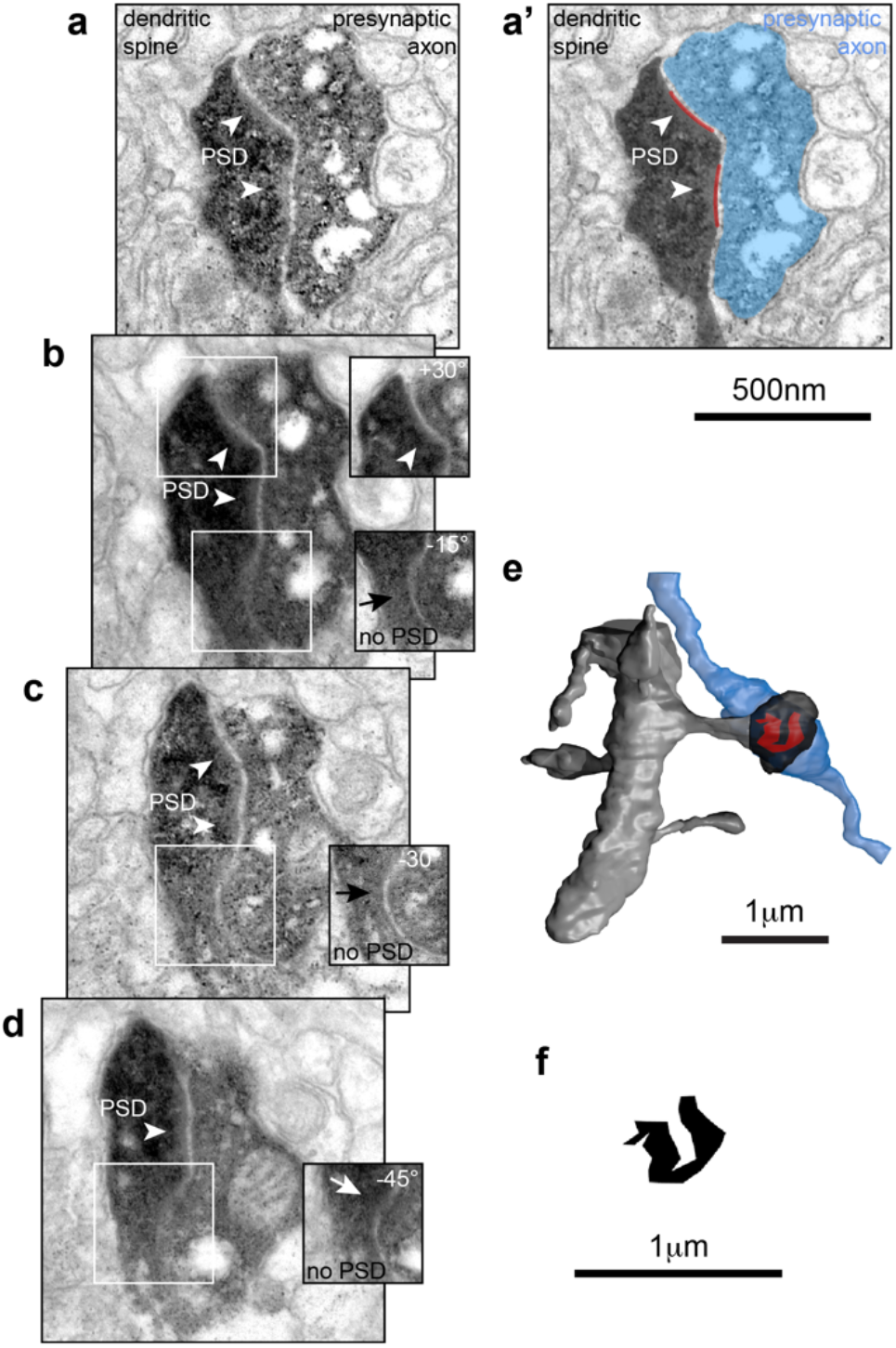
Reconstruction of PSD area in biocytin-filled dendritic spines. **a - d** Consecutive series of electron micrographs through a synapse that was previously recorded from (synapse 3 in Fig. 1, micrographs rotated). All sections acquired without tilt angle; white boxes, regions in which imaging plane was not perpendicular to synaptic cleft and PSD identification was hindered; insets, tilted micrographs of these regions (tilt angle indicated). In this experiment, the PSD appeared as a negative staining against biocytin reaction product. Arrowheads, PSD; arrows, locations where no PSD was identified. **a’** Same micrograph as in **a** with synaptic structures annotated for 3D EM reconstructions; blue, presynaptic axon; black, postsynaptic dendrite; red, PSD. **e** Complete 3D EM reconstruction of the entire synaptic contact; blue, axon; grey, dendrite; black, spine head; red, PSD. **f** *En-face* representation (2D projection) of the reconstructed PSD reveals a complex ‘horseshoe-like’ morphology.

**Extended figure 2.**
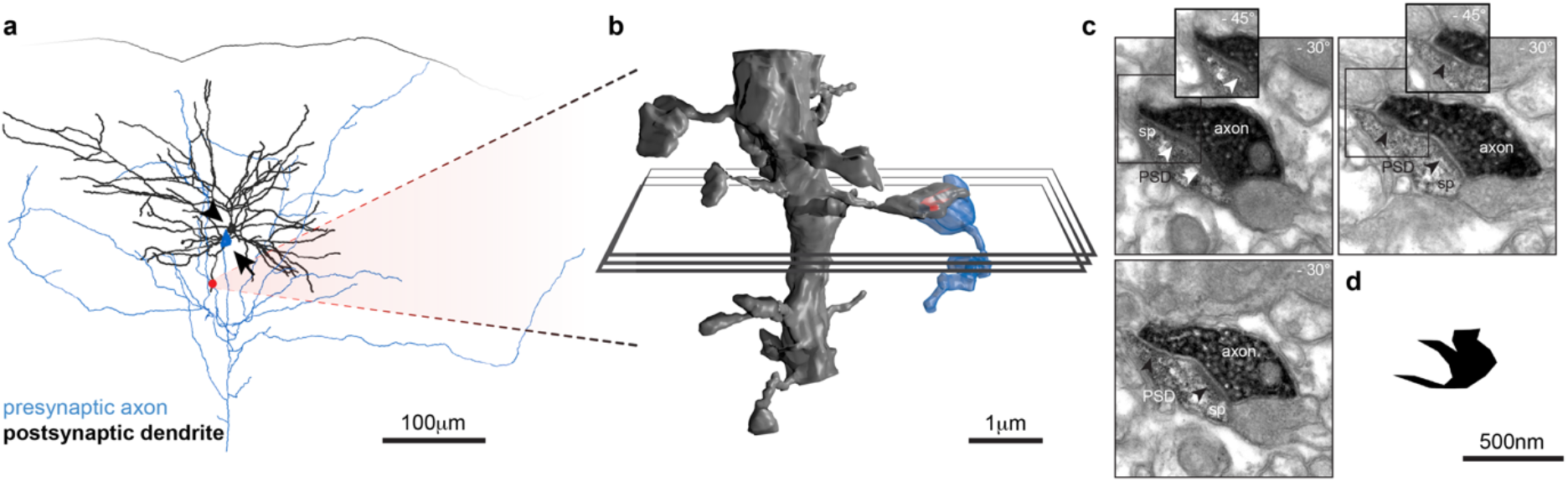
High-fidelity correlated LM–EM of biocytin-labeled synapses following paired recordings. A single synapse was formed between connected pyramidal neurons. **a** Full 3D LM reconstruction of presynaptic axon and postsynaptic dendrite of synaptically connected biocytin-stained pair; a single axodendritic apposition was identified between the cells in LM (red dot); postsynaptic axon and presynaptic dendrite excluded for clarity; arrow, soma of presynaptic neuron; arrowhead, soma of postsynaptic neuron; cortical surface indicated. **b** 3D EM reconstruction of the apposition; same color scheme as in a; red, PSD; planes indicate z-positions of electron micrographs in c. **c** Electron micrographs of consecutive cross-sections through the synapse. The dendritic spine (sp) was clearly stained, while biocytin appeared patchier than in other experiments, allowing for easy PSD identification (arrowheads); micrographs were acquired at a tilted angle along the axis of the synaptic cleft for a clear view on the PSD (tilt angles indicated); insets, additional tilts of regions surrounded by boxes. **d** *En-face* representation (2D-projection) of reconstructed PSD.

**Extended figure 3.**
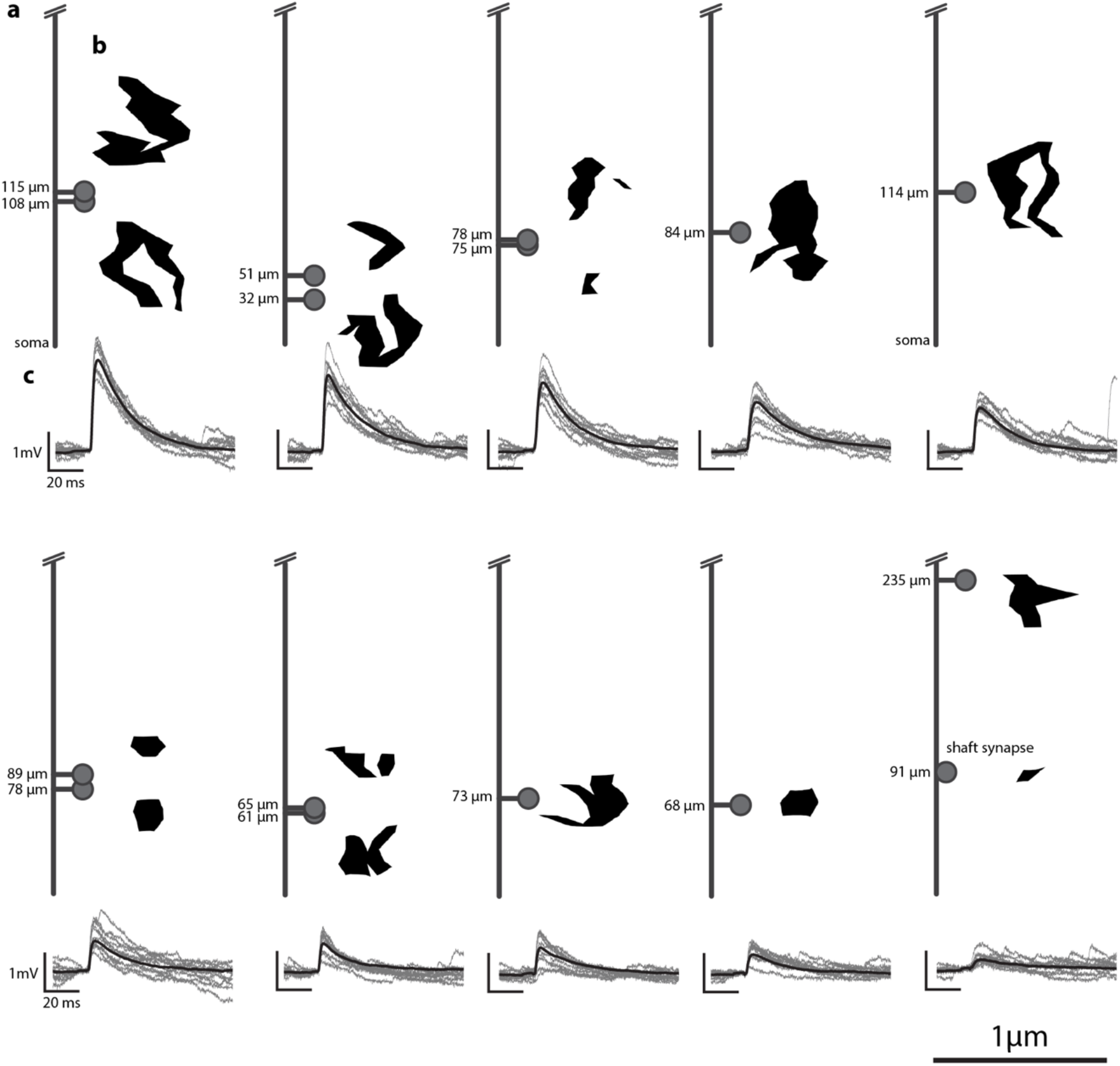
Overview across all 10 experiments showing for each connection the important anatomical properties of recovered synapses and corresponding evoked EPSP waveforms. Experiments sorted by decreasing mean EPSP amplitude. **a** Dendritic distances of the identified synapses to the soma. The collapsed dendritic tree is represented schematically ranging from soma (bottom) to a distance of 250 μm (top, cut off); dendritic distances at which respective synapses were found indicated by spine locations. No distinction made between apical and basal dendrites; all synapses were formed on dendritic spines, except of the proximal synapse in experiment 10, which was formed on dendritic shaft (indicated). **b** Morphologies of reconstructed PSDs. *En-face* representations (2D-projections) of PSDs highlight the ranges of identified sizes and shapes; PSDs were positioned next to spines according to their respective dendritic distances as indicated in a; corresponding scale bar indicated at bottom right of figure. **c** Waveforms of evoked EPSP recordings; black, average waveform of EPSP recordings during stable epochs for each connection; grey, 10 randomly-selected single-trial EPSP waveforms taken during stable epochs of recording.

**Extended figure 4.**
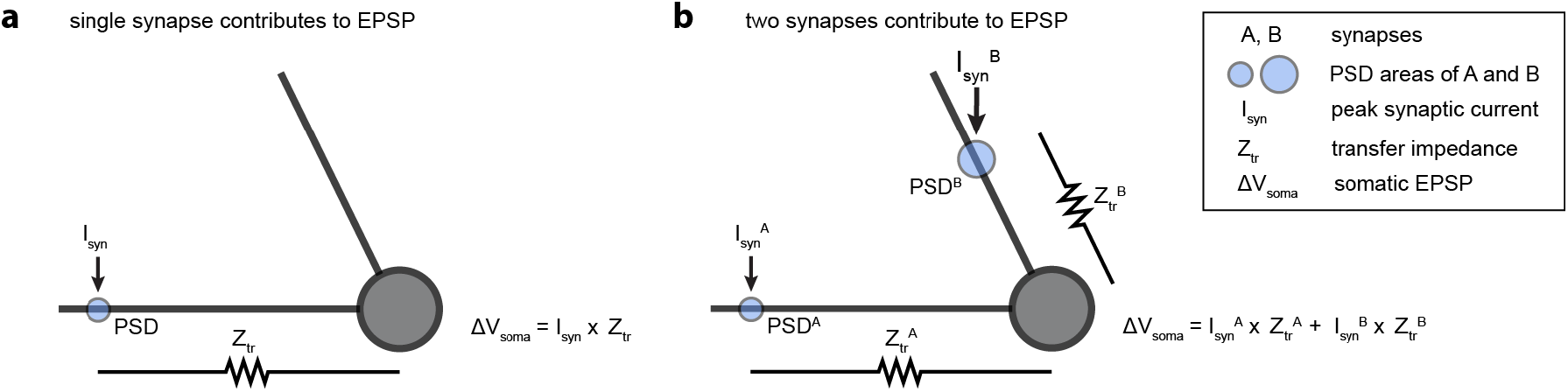
Approximating peak synaptic current from EPSP amplitude and transfer impedance. **a** For connections comprised of a single synapse, peak synaptic current can be estimated following Ohm’s Law from the experimentally recorded somatic EPSP amplitude and the simulated distance-dependent transfer impedance. **b** When two synapses (A, B) were identified in EM, we additionally assumed that the ratio of I_syn_^A^ to I_syn_^B^ followed the ratio of PSD^A^ to PSD^B^ and that synaptic currents summate linearly at the soma.

**Extended figure 5.**
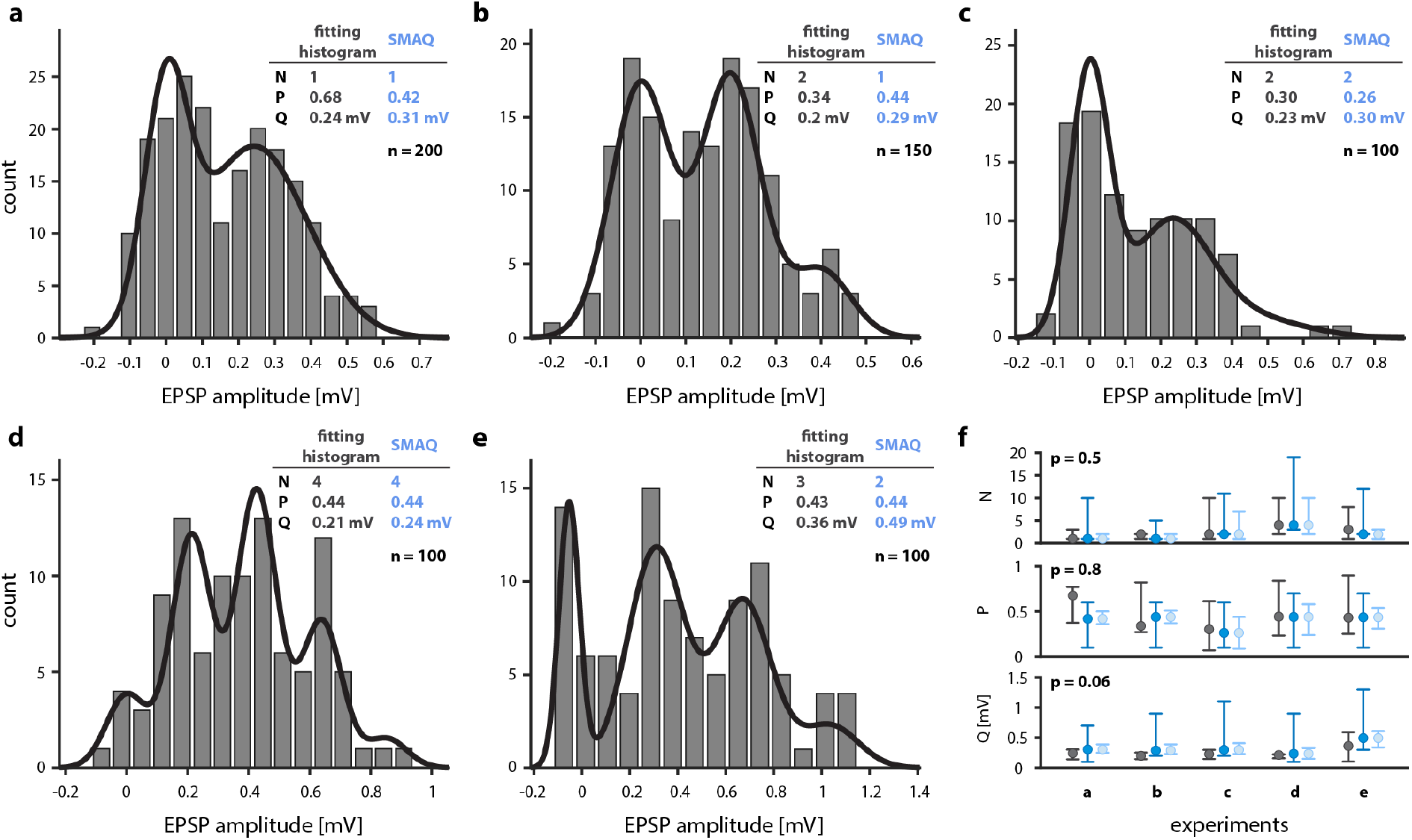
Quantal analysis using SMAQ provides similar results and uncertainty as the method of fitting binomial models to peaky histograms. **a – e** Five EPSP amplitude histograms contained equally-spaced peaks. These histograms could be fit successfully with a simple quantal binomial model to extract quantal parameters (black lines overlaid on histograms) and in addition, quantal parameters were estimated using SMAQ; insets, comparisons of the quantal parameters found by fitting peaky histograms and SMAQ; N, number of release sites; P, release probability; Q, quantal size; n, number of entries per histogram. **f** Comparison of solutions for quantal parameters and associated 95% confidence intervals given by the method of fitting peaky histograms (black) and SMAQ (blue) across the five experiments shown in **a-e**. Dark blue, Bayesian-inspired confidence intervals for SMAQ; light blue, confidence intervals for SMAQ derived from the same bootstrap resampling algorithm used to derive confidence intervals for the method of fitting peaky histograms (black). Solutions provided by the two methods were compared with the non-parametric Wilcoxon matched-pairs test (two-tailed p-value reported). For all three quantal parameters, solutions given by the two methods were not significantly different (see text for details).

## Methods

### Reagents

All solutions were prepared in MilliQ water (18.2 MOhm cm). Chemicals for electrophysiology solutions were purchased from Sigma-Aldrich. Chemicals for histology, including phosphate buffer (PB), phosphate buffered saline (PBS), and tris-buffered saline (TBS) were made in-house.

### Animals

Male B6/C57 mice between postnatal days 21 to 40 under the license of K.A.C.M. were used in the study. Handling and experimental procedures were approved by the Cantonal Veterinary Office, Zurich, Switzerland.

### Slicing

Animals were anesthetized with isoflurane and quickly decapitated. After removal of the brain, 300 μm thick sections were quickly sliced in ice-cold artificial cerebrospinal fluid (ACSF; containing, in mM, 87 NaCl, 75 sucrose, 26 NaHCO_3_, 2.5 KCl, 1 NaH_2_PO_4_, 0.5 CaCl_2_, 7 MgSO_4_, 10 glucose and continuously oxygenated with 95% O_2_, 5% CO_2_). A para-coronal slicing angle was used to preserve apical dendrites parallel to the cutting plane, which enhanced the probability of finding connected pairs. Slices were allowed to recover for 30 min in oxygenated recording ACSF at 36 °C (containing, in mM, 119 NaCl, 2.5 KCl, 26 NaHCO_3_, 1.25 NaH_2_PO_4_, 1.3 MgSO_4_, 2.5 CaCl_2_, and 10 glucose, constantly perfused with 95% O_2_, 5% CO_2_). The solution containing the slices was then kept at room temperature until the recordings.

### Electrophysiology

A crucial technical necessity of the study was to recover axons and dendrites of recorded neurons completely and prevent structural damage to enable high-fidelity ultrastructure for subsequent EM. We achieved this by optimizing intricately the composition of pipette solution and the shape of recording pipettes. The pipette solution contained, in mM: 115 K-gluconate, 20 KCl, 2 Mg-ATP, 2 Na-ATP, 10 Na-phosphocreatin, 0.3 GTP, 10 Hepes, pH was set to 7.2 with KOH. In a subset of experiments, K-gluconate was decreased to 105 mM to enable increased biocytin concentrations (~ 1%), which were found to enhance greatly complete fillings. No significant differences in input resistance (R_in_) and membrane time-constants (tau_m_) were found between the two recording conditions. Pipette solution osmolarity including biocytin was adjusted to 290–300 mOsm, which slightly exceeded the osmolarity of the recording ACSF and greatly reduced rapid swelling of dendrites while allowing for complete diffusion of biocytin throughout fine neurites. Patch pipettes were pulled from borosilicate capillaries (1.5 mm outer diameter, 1.17 mm inner diameter, Warner Instruments) using a P-97 Flaming/Brown micropipette puller (Sutter Instrument). Pipettes with elongated tapers significantly reduced swellings of dendrites. Pipette tip diameters ranged between 2-3 μm and pipette resistance was between 6 and 8 MOhm. Slices were placed in a submersion chamber and constantly perfused with warmed, oxygenated ACSF (2-3 ml/min), the temperature at the center of the chamber was maintained at 33–35 °C. Cells were visualized under an upright microscope (Olympus BX61W1) equipped with infrared differential-interference contrast (IR-DIC) optics and 10x and 60x water-immersion objectives. Whole-cell somatic patch-clamp recordings were established from pyramidal neurons in L2/3 of mouse barrel cortex in current-clamp mode (Multiclamp-700A amplifier, Axon Instruments). Data was sampled at 10 kHz, filtered at 3 kHz, and digitized using an AD converter (Digidata 1322, Axon Instruments). Recordings were visualized and controlled using the pClamp software (Molecular Devices). After establishing the whole-cell configuration, access resistances (R_access_) was measured, which typically ranged from 15 to 25 MOhm. Recordings with R_access_ > 30 MOhm were discarded. Bridge potential was compensated for and liquid-junction potential was not corrected for. The resting membrane potential (V_m_) immediately after establishing whole-cell access ranged between −85 to −75 mV. De- and hyperpolarizing current pulses were injected to measure R_in_ and tau_m_ from the current-voltage (I-V) traces. We simultaneously recorded pairs of closely proximate L2/3 pyramidal neurons and tested for synaptic connections by evoking single action potentials alternatingly in the two cells at 0.2 Hz. We identified connections by averaging 20 – 50 trials and searching for evoked EPSPs immediately following action potential firing. Once a synaptic connection was identified, single action potentials were evoked in the presynaptic neuron at 0.1 Hz or 0.2 Hz. If necessary, we used a holding current to maintain V_m_ of the postsynaptic neuron below −70 mV to ensure that EPSPs were only mediated by AMPAr currents^23,41,42^; however, this was rarely needed. To control that evoked EPSPs were AMPAr mediated, the decay phases of averaged EPSP waveforms were fit with single exponentials, which was successful in all experiments and confirmed that NMDAr (N-methyl-D-aspartate receptor) currents did not contribute significantly to our EPSPs recorded below −70 mV^50^. Evoked EPSPs were recorded for up to two hours or for as long as the preparation remained stable.

### Measurement of EPSP amplitudes

The peak amplitude of each recorded EPSP was measured offline using a custom-written Matlab package as in^23,41,42^. Briefly, V_m_ was computed by averaging the membrane voltage in a 1.5 - 2 ms time window before EPSP onset and subtracted from a measurement of the peak EPSP using the same window over the EPSP peak. Window width was chosen to include EPSP peak and exclude rise and decay phase. Spacing of baseline and peak windows was chosen so the baseline potential was measured as closely to EPSP onset as possible to minimize noise in EPSP recordings. Typical window spacings ranged from 5 to 6 ms (range: 4.5 – 7 ms). Window width and spacing remained constant within experiments. To obtain an independent measure of noise associated with V_m_, a separate set of identical windows was used on a portion of the membrane potential preceding the evoked EPSP.

### Selection of EPSP data

For reliable measurements of mean EPSP amplitude and subsequent quantal analysis, we required extended periods of stable recordings. We imposed stability criteria on our data and analyzed only a single epoch of EPSP recording that remained stable for at least 100 consecutive trials. Stable epochs were defined as those in which mean and standard deviation of evoked EPSPs, as measured in blocks of 25 trials, remained close to their values in a reference block. The mean was required to remain within 3x the standard error of the mean and the standard deviation was not allowed to change by more than 30%^23,41,42^. Wherever possible, we sought to analyze the earliest epochs of recording after pipette break-in. Therefore, when using the initial block of recording as reference block yielded a stable epoch of ≥ 100 consecutive trials, this epoch was analyzed. Otherwise, we iteratively assigned the remaining blocks as reference blocks and computed the stable epoch for each of them. The longest stable epoch was then chosen for analysis. Therefore, different epochs of EPSP recordings after pipette break-in were used between experiments and we sought to exclude the possibility that different degrees of dialysis with pipette solution and resulting cytosol washout had compromised our dataset^23^. Thus, we related cumulative PSD area with the mean EPSP amplitude from only the initial 25 trials of recording for all experiments. Reassuringly, this revealed the same correlation efficient (r = 0.73, p = 0.02) as the correlation between cumulative PSD area and mean EPSP amplitude calculated from the ≥ 100 consecutive trials (Fig. 3 d).

### Measurements of EPSP kinetics

50 – 100 trials of stable epochs of recordings, in which EPSPs were evoked, were aligned to presynaptic action potential peak, Vm was subtracted, and they were averaged. Measurements were performed on averaged traces in Stimfit^51^. EPSP rise-time was calculated as the interval between 20% and 80% of EPSP peak amplitude. Exponentials were fit to the EPSP decay phase using Clampfit 9 (Axon Instruments).

### Histology

Immediately following recordings, slices were fixed overnight in 4% paraformaldehyde (PFA), 0.5% glutaraldehyde, and 15% picric acid in 0.1 M PB, pH 7.4. Slices were then washed in PB, incubated in an increasing sucrose ladder for cryoprotection, and rapidly frozen in liquid nitrogen. To quench endogenous peroxidases, sections were incubated in 10% methanol, 3% hydrogen peroxide (H2O2) in PBS. After washing, sections were reacted with the Vectastain ABC Kit (Vector Laboratories, catalog # PK-6100, RRID: AB_2336819) overnight at 4 °C. After washing, biocytin was visualized with a protocol containing nickel-diaminobenzidine (Ni-DAB) tetrahydrochloride and H2O2 treatment. The reaction was terminated with a series of washes in PB.

### Re-slicing and embedding

To allow for complete reconstructions of recorded neuron pairs in LM, sections were re-sliced to 80 μm. Briefly, sections containing completely filled neurons, as assessed by LM after the Ni-DAB reaction, were carefully glued flat onto a block of agar using UHU superglue gel (UHU GmbH & Co. KG). The slice surface that was recorded from pointed upwards. Slices were immediately embedded with lukewarm agar that was allowed to solidify at 4 °C. The block was trimmed and placed under a vibratome in PB. The embedding agar provided the necessary stability to carefully re-slice sections to 80 μm and completely prevented tissue loss in the process. Thin sections were carefully collected in PB and treated in 1% osmium tetroxide in PB for 10 – 20 min, depending on section thickness and reaction speed. Sections were dehydrated using an ascending series of ethanol and propylene oxide (including treatment in 1% uranyl-acetate in 70% ethanol), and flat-mounted in Durcupan resin (Sigma-Aldrich).

### LM reconstructions

Neuron pairs were included only when presynaptic axon and postsynaptic dendrite were completely filled with biocytin and could be entirely reconstructed in LM. To assess completeness of filling, we examined whether staining of neurites faded out within sections, which led us to immediately discard experiments. Only when all respective neurites terminated as low or high endings at the surfaces of the original 300 μm section, or ended in well-labeled terminals, these neuron pairs were used in the study. 3D morphologies of pre- and postsynaptic neuron were fully reconstructed in the Neurolucida Software package (MicroBrighField) under an Olympus BX 51 light microscope equipped with a 60x and 100x oil objective. We identified and marked all appositions between presynaptic axon and postsynaptic dendrite in LM. Morphological criteria for determining a contact included the existence of an axonal bouton and no discernable gap between axon and postsynaptic dendrite or spine. Axons traversing within 4 μm above or below a postsynaptic dendrite and forming a bouton close to the crossing point were also marked as contacts and subjected to EM. In principle, a dendritic spine could extend in the z-direction towards the axon, which could remain disguised by the traversing axon. In fact, we discovered synapses at two such contacts. In the rare cases when a contact was obscured by leakage of biocytin from labeled dendrites, we completely reconstructed the respective contact in EM, which allowed us to verify the existence of synapses at these sites unambiguously in all cases. LM reconstructions were exported into the Blender software and the dendritic distance to soma was measured for all contacts using the NeuroMorph Plugin.

### Correlated LM-EM

Tissue blocks containing LM appositions were serially sectioned at 60 nm and collected on pioloform-coated single-slot copper grids. Low-magnification electron micrographs were correlated with LM overview images of the same region taken before ultrathin sectioning using the TrakEM2 plugin of ImageJ. Comparing blood vessel patterns and labeled neurites across the neuropil allowed us to localize and recover all LM appositions in EM. To verify whether LM appositions were synapses, serial electron micrographs (13,500x) were generated for all appositions and loaded into TrakEM2 for reconstruction. When synapses were found, series of high-magnification images (46,000x) were acquired for the entire synapse for high-magnification reconstructions. We have not corrected for tissue shrinkage through the histology procedures in any of our quantitative EM measurements. We have measured shrinkage throughout all stages of processing in both cat^52^ and mouse neocortex (unpublished observations). Aldehyde fixation–perfusion produced a consistent 11% shrinkage.

### Synapse identification and reconstruction in EM

Synapses between biocytin-filled presynaptic axon and postsynaptic dendrite were identified from series of electron micrographs as follows: they were required to possess a vesicle-filled presynaptic bouton, form axodendritic contacts in multiple consecutive sections separated by a synaptic cleft, and contain a PSD in the dendrite. Identification of synaptic vesicles and synaptic cleft was largely unaffected by biocytin staining. To ensure that we could reliably reconstruct and measure PSD area, we implemented four additional procedures. First, we tilted the ultrathin sections containing synapses along the dimension of the synaptic cleft at six angles in the EM (−45 °, −30 °, −15 °, +15 °, +30 °, +45 °), which allowed us to obtain an optimal perpendicular imaging plane through the synaptic cleft for all sections. Second, colors of micrographs were inverted, which highlighted subtle contrast differences and aided the identification of the PSD. Third, each synapse was reconstructed by two experts independently and in a blinded manner with respect to the physiological features of synaptic transmission and a consensus reconstruction was found between the two reconstructions. Fourth, we reconstructed a large number of unlabeled PSDs and their dendritic spine heads (n = 75) from adjacent neuropil. Unlabeled synapses were randomly selected as described elsewhere^35^. We then compared the identified PSD-spine head relationship of the biocytin-labeled synapses and the unlabeled synapses from the same neuropil using Fisher-Z-transformation. 3D reconstructions of representative structures were exported into the Blender software, fitted with a skin, and rendered to offer a 3D impression.

### Statistical robustness of cumulative PSD – EPSP correlation

For each experiment, 100 resampled EPSP distributions were generated by drawing with replacement from the recorded EPSP distributions. The same number of samples was drawn that was present in the original distributions. A small amount of jitter was added to each selected EPSP as a random number drawn from a Gaussian process with mean of 0 mV and a standard deviation of ¼ of the standard deviation of the recording noise of the respective experiment^23^. Then, the mean EPSP amplitude was computed for each resampled distribution. We generated new scatter plots of cumulative PSD area versus mean EPSP amplitude by randomly selecting a mean EPSP amplitude from the 100 resampled distributions per experiment. The Spearman correlation coefficient was computed for 10,000 scatter plots that were randomly picked from a possible 10^11^ combinations. To compare the robustness of the correlation on a trial-to-trial basis, we generated scatter plots by drawing with replacement a single EPSP amplitude for each experiment. Likewise, we computed the Spearman correlation coefficient for 10,000 randomly-drawn correlations.

### Compartmental NEURON model

A compartmental model was generated in the NEURON software package from a volumetric reconstruction of a L2/3 pyramidal neuron using the d-lambda rule. We chose a cell, whose anatomical and physiological properties best represented the postsynaptic neurons in the study (mean ± standard deviation of population in brackets). This cell had a total dendrite length of 5285 μm (4611 ± 1440 μm), longest dendrite of 842 μm (857 ± 238 μm), R_in_ of 72.9 MOhm (67 ± 12 MOhm), and tau_m_ of 11.8 ms (14.8 ± 3.9 ms). The experimentally recorded waveforms to two hyperpolarizing and one depolarizing current step (−40 pA, −80 pA, +40 pA) were used to fit the specific membrane resistance (R_m_ = 5245.2 Ohm cm^2^), specific membrane capacitance (C_m_ = 4.26 μF/cm^2^), and specific axial resistance (R_a_ = 114.3 Ohm cm) of the model. These values reflect that spines were not explicitly modeled. V_m_ was set to the experimentally observed −82.6 mV. After tuning the model, R_in_ (72.7 MOhm) matched the experimental R_in_ (72.9 MOhm). Alpha synaptic conductances (g_max_ = 2 nS, tau_max_ = 0.7 ms) were simulated at 810 different locations on the dendritic tree. Synapses simulated at distances of 40 to 120 μm from the soma (i.e. within the distance range of synapses found in our experiments) evoked somatic EPSPs with comparable kinetics as experimentally recorded EPSPs (peak amplitude: 0.8 ± 0.2 mV and 1.1 ± 0.7 mV, 20% - 80% rise time: 1.3 ± 0.4 ms and 1.4 ± 0.3 ms, time-to-peak: 3.9 ± 1.4 ms and 3.9 ± 0.8 ms, decay time constant: 19.7 ± 3.2 ms and 18.6 ± 2.7 ms, respectively; mean ± standard deviation). Transfer impedance (Z_tr_) was calculated as somatic EPSP amplitude divided by peak synaptic current (I_syn_). Because I_syn_ is negative by convention, all values of Z_tr_ were given a positive sign. Additionally, we computed Z_tr_ using the impedance tool of the NEURON simulation environment at a frequency of 100 Hz (Z_tr_^100 Hz^). The resulting distributions of Z_tr_ and Z_tr_^100 Hz^ matched well and were statistically not different (p = 0.62, non-parametric Mann-Whitney test, two-tailed p value). Z_tr_ was plotted as a function of synaptic distance to soma and a double exponential function was fit to the data. To quantify uncertainty of the model associated with Z_tr_, dendritic distances were binned (into 10 μm windows or larger windows, which included at least 20 entries) and 2.5 and 97.5 percentiles of Z_tr_ calculated for each bin.

### Approximation of peak synaptic current

We computed cumulative peak synaptic current (cI_syn_) for our 10 connections by dividing the EPSP amplitude by Z_tr_ (as derived from the fit) that corresponded to the mean dendritic distance of the synapses made by the connection. We approximated I_syn_ for each of the 16 synapses separately, as follows. When a single synapse was found for a connection, I_syn_ was approximated by dividing the EPSP amplitude by Z_tr_ for the respective synaptic distance from soma as derived from the fit. 95% confidence intervals were constructed from the 2.5 and 97.5 percentiles of Z_tr_ in the respective distance bin. When connections contained two synapses (A and B), two additional assumptions were necessary to estimate I_syn_^A^ and I_syn_^B^. Motivated by the cumulative PSD-cI_syn_ correlation, we assumed that the ratio **R** of PSD areas (PSD^A^ to PSD^B^) is reflected in the ratio of I_syn_^A^ to I_syn_^B^. Note that while this assumption constrains the ratio of I_syn_^A^ to I_syn_^B^ within an experiment, importantly, it does not bias the comparison of absolute I_syn_^A^ and I_syn_^B^ between experiments. Also, it does not imply that the two attenuated EPSPs at the soma follow the same ratio. Because of different synaptic distances from soma, they will be attenuated by different degrees. Finally, we assumed that the resulting attenuated EPSPs summed linearly at the soma to generate the EPSP amplitude^36^. Using these assumptions, the respective I_syn_ was calculated for each of the two synapses, given the EPSP amplitude, Z_tr_ for each synapse (Z_tr_^A^ and Z_tr_^B^), and **R**:

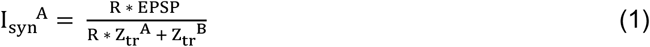

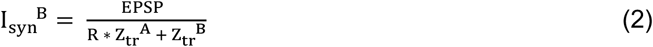

As above, 95% confidence intervals were constructed for I_syn_^A^ and I_syn_^B^ by using the values for the 2.5 and 97.5 percentile of Z_tr_^A^ and Z_tr_^B^ in the respective distance bins.

### Statistical Moments Analysis of Quanta

The binomial equations for mean (μ), standard deviation (σ), and skewness (γ) of experimentally recorded EPSP distributions,

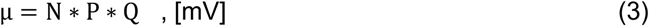

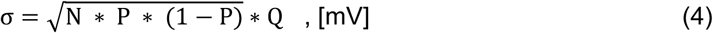

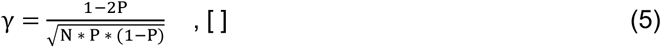

were reshaped to derive unique analytical solutions for the number of release sites (N), release probability (P), and quantal size (Q), as functions of μ, σ, and γ:

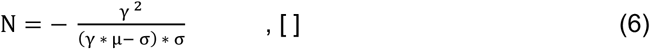

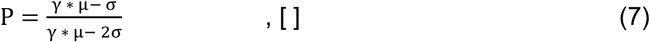

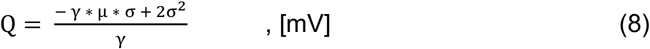

For each connection, μ, σ, and γ were computed for the stable epoch of EPSP recording, and the corresponding N, P, Q calculated. Inspired by Bayesian logic, we derived 95 % confidence intervals for the quantal parameters by asking: which underlying binomial models (combinations of N, P, Q) could have generated EPSP distributions that could have provided the same SMAQ solutions? We simulated binomial models from all possible permutations of a large range of quantal parameters (N between 1 and 20, P between 0.1 and 0.9, Q between 0.1 mV and 1.5 mV, n = 2700). We generated individually-tailored confidence intervals for each one of our experiments: 10,000 realizations were simulated for each of the 2700 models, which contained the same number of trials as the experimentally observed histogram. Additionally, we added the experimentally observed recording noise onto each simulated EPSP as a randomly drawn realization from a Gaussian with a mean of zero and the same standard deviation as measured for V_m_ (i.e. the recording noise) in experiment. We then used SMAQ to calculate N, P, and Q for each one of these 27 * 10^6^ simulated EPSP distributions and asked: which underlying binomial models could have produced the experimentally observed N, P, and Q at the 95% certainty level? For example, we generated the distribution of N of all models that had produced the same SMAQ solution for N as the experimental histogram. The 2.5 and 97.5 percentiles of that distribution gave the 95% confidence intervals of the experimentally observed N and indicated which underlying Ns, with 95% certainty, could have also produced the experimentally observed EPSP distribution. Additionally, we computed 95% confidence intervals using the same bootstrap resampling algorithm, which is used to create confidence intervals for quantal parameters derived by fitting binomial models to peaky histograms (see below). This allowed us to compare the uncertainties associated with the solutions of SMAQ and the method of fitting binomial models to peaky histograms.

### Fitting binomial models to peaky histograms

Stable EPSP amplitude histograms that revealed equally-spaced peaks were fit with a quantal binomial model as explained in^23,41,42^. We used the MATLAB implementation of the method available on *www.jennyreadresearch.com*. The model was allowed to search for an optimal N in the range of 1 to 20. Noise was not constrained to allow for negative quantal variance^53,54^, which was observed in one experiment, and is otherwise not implemented in the method. To enable a fair comparison with SMAQ, conductance probability was set to 1 and offset was disabled unless no successful fit could be found. Only in one case, an offset of −0.053 mV had to be implemented. All seven available adequacy-of-fit tests were used, which include the Kolmogorov–Smirnov D statistic, the sum of squared differences between model and data cumulative distributions, and chi-squared statistics for five different bin sizes. A fit was considered successful only when it passed all tests. We used the inbuilt bootstrap-resampling function to calculate 95% confidence intervals for the best-fit parameters. Briefly, new distributions were generated by drawing with replacement from the experimental EPSP histogram. A small amount of jitter was added to each selected EPSP and the resampled EPSP distributions were fit and tested for adequacy in the same manner as the experimental distribution. The procedure was repeated until 100 successful resampled fits had been generated. 95% confidence intervals were constructed for N, P, and Q from the 100 estimates of N, P, and Q of the resampled distributions.

### Statistics

Non-parametric tests were used for all statistical analyses. A significance level of 0.05 was used. Statistical tests were performed in Prism 6 (GraphPad). Correlation analyses were performed with a two-tailed nonparametric Spearman test. Lines were fit through scatter plots with linear regression.

## Acknowledgements

We are grateful to Ora Ohana for her training and sharing her expertise in patch clamp recordings. We thank Sergio Solinas for assistance with the NEURON simulation and Saray Soldado-Magraner and John C Anderson for their support. We thank Andrew Bolton, Moritz Buchholz, and Florian Engert for reading the manuscript. As members of the Institute of Neuroinformatics, the authors are signatories of the Basel Declaration. This work was supported by the Swiss Society for Neuroscience, the Neuroscience Center Zurich, and the Swiss National Science Foundation (G.F.P.S.).

## Author Contributions

Authors are listed alphabetically.

K.A.C.M. and G.F.P.S. designed experiments;

G.F.P.S. performed electrophysiology experiments;

S.H.-R., G.K. and G.F.P.S. performed histology, LM reconstructions, and correlated LM-EM;

G.K. and G.F.P.S. performed PSD reconstructions;

G.F.P.S. and K.J.S. analyzed electrophysiology data;

G.F.P.S., and K.J.S. developed SMAQ;

K.A.C.M. and K.J.S. supervised the work;

K.A.C.M., G.F.P.S., and K.J.S. wrote the paper.

